# Endolysosomal damage promotes intraluminal protein condensate formation that limits cytosolic leakage

**DOI:** 10.64898/2026.05.25.726725

**Authors:** Xu Li, Yu Zhang, Kaiqiang Zhao, Gu Chen, Di Cui, Mehran Nikan, Stephen G. Young, C. Frank Bennett, Punit P. Seth, Tao Ni, Maximiliano G. Gutierrez, Haibo Jiang

**Affiliations:** Department of Chemistry, The University of Hong Kong, Pok Fu Lam, Hong Kong, China; School of Biomedical Sciences, Li Ka Shing Faculty of Medicine, The University of Hong Kong, Hong Kong, China; Ionis Pharmaceuticals, Inc., Carlsbad, CA 92010, USA; Departments of Medicine and Human Genetics, University of California, Los Angeles, CA 90095, USA; Alnylam Pharmaceuticals, Inc., Cambridge, Massachusetts 02142, USA; The Francis Crick Institute, London, UK; Laboratory for Synthetic Chemistry and Chemical Biology Limited, Health@InnoHK, Innovation and Technology Commission, Hong Kong, China

## Abstract

Endolysosomal membrane damage is a detrimental process in mammalian cells that results in leakage of the luminal contents into the cytosol. However, the nature and extent of the leakage during membrane damage is unknown. Here, we show that endomembrane damage induces the rapid formation of intraluminal condensates in endolysosomes. A subset of resident luminal proteins undergo spatially coordinated condensation upon endomembrane damage. Electron microscopy reveals distinct luminal morphology, and cryo-electron tomography confirms the condensed ultrastructure in their native state. Condensate formation occurs across mechanistically distinct modes of membrane injury and is reversibly dissociated upon lysosomal recovery. Remarkably, these condensates impose a previously unrecognised barrier to endolysosomal escape of therapeutic oligonucleotides. Despite endomembrane damage, luminal oligonucleotide therapeutics are sequestered in damaged endolysosomes through condensate-mediated biophysical immobilisation. Targeting condensate sequestration could represent a novel strategy to improve oligonucleotide-based therapeutics.

---

Endolysosomal damage is a threat to cellular health, arising from pathogens, crystals, and membrane-permeating compounds^1–4^. Recent studies have uncovered a repertoire of repair mechanisms that respond to endomembrane damage, including the ESCRT machinery for small membrane lesions^5,6^, the phosphoinositide signalling pathway for ER-lysosome lipid exchange^7^, annexins for larger wounds^8,9^, and stress granules that have been proposed to stabilize ruptured endomembrane^10^. These responses are fundamentally focused on restoring the integrity of the limiting membrane. On the other hand, very little is known about the leakage of the luminal contents during damage and repair. The prevailing view has largely assumed that luminal cargo when released to the cytosol follow a passive leakage through membrane pores in a damage size-dependent manner^11–14^. Yet emerging evidence suggests that the endolysosomal lumen is not a simple aqueous compartment but instead is a highly complex environment that sensitive to changes in pH and ionic conditions^15–18^. Whether this sensitivity enables an compartment-intrinsic response that influences the behaviour of luminal contents upon membrane injury is unknown.

Physiological contexts for limited, non-lethal endolysosomal leakage have been described across diverse cell types and processes, including host defence^19^, antigen cross-presentation^12,20,21^, and mitotic cell division^22^, where controlled endolysosomal damage has been proposed to facilitate chromosome segregation. Constitutive pores on neuronal endolysosomes further indicate that the endolysosomal membrane need not be intact for cellular function^23^. These studies suggest that luminal contents are not unrestrictedly released in a size-dependent membrane damage manner. However, whether there are cellular mechanisms that selectively retain luminal contents in damaged endolysosomes is unclear.

Engineering the escape of oligonucleotide therapeutics from endosomes is a major challenge in drug development. The vast majority of internalized oligonucleotides become associated within the endolysosomal system^24–26^. Even when lipid nanoparticle-encapsulated oligonucleotides are used to induce endolysosome damage and leakage^27–30^, dosing regimens typically involve hundreds of times more oligonucleotides than theoretically required^31^. Chemical modifications and ligand-conjugation strategies (*e*.*g*., GalNAc and GLP1-peptide) have improved nuclease resistance^32,33^ and cell type-specific uptake^34,35^. Despite these advances, however, endosomal escape efficiency remains the principal bottleneck limiting broad application of oligonucleotide-based therapeutics^29^. Currently, we do not understand the mechanisms governing endosomal escape of oligonucleotide-based therapeutics.

Here, using super-resolution live-cell imaging, electron microscopy, and cryo-electron tomography, we show that endomembrane injury triggers the rapid formation of intraluminal condensates. Condensate formation and dissociation are coupled to the state of endolysosomal homeostasis. We further show that these condensates sequester luminal antisense oligonucleotides (ASOs) by immobilization, suggesting an unrecognized, condensate-mediated barrier to endosomal escape. Together, our findings reveal that the endolysosomal lumen is not a passive bystander during membrane damage but an active player that regulates the selective leakage of endolysosomal cargo.

## Results

### Intraluminal condensates are rapidly formed after endolysosomal damage

We first studied the behaviour of luminal contents upon L-leucyl-L-leucine methyl ester (LLOMe) treatment, which destabilises endolysosomal membranes by cathepsin-mediated processing^36^. Live-cell imaging of U2OS cells expressing mNeonGreen-LAMP1 and loaded with 10-kDa Alexa Fluor 647–dextran revealed that LLOMe triggered a rapid morphological change within the lumen of endolysosomes. Under steady-state conditions, fluid-phase dextran is localised in endolysosomes. Strikingly, upon LLOMe treatment, dextran underwent a dramatic redistribution, condensing into intraluminal bright puncta (**Fig. 1a** and **Video S1-S4**). Quantitative analysis of these events showed a sharp increase in fluorescence within individual endolysosomes. This increase in intensity was accompanied with a rise in maximal intensity with the mean intensity remained stable, suggesting condensate formation (**Fig. 1b**). To determine whether the condensation was a general response rather than an rare event observed in selected endolysosomes, we quantified the mean of maximal intensity of all individual dextran puncta per cell before and after membrane damage. This analysis confirmed that LLOMe treatment induced a significant increase in the localized maximal intensity for both 10 kDa and 70 kDa dextran cargo (**Fig. 1c, d**).

**Figure 1.**
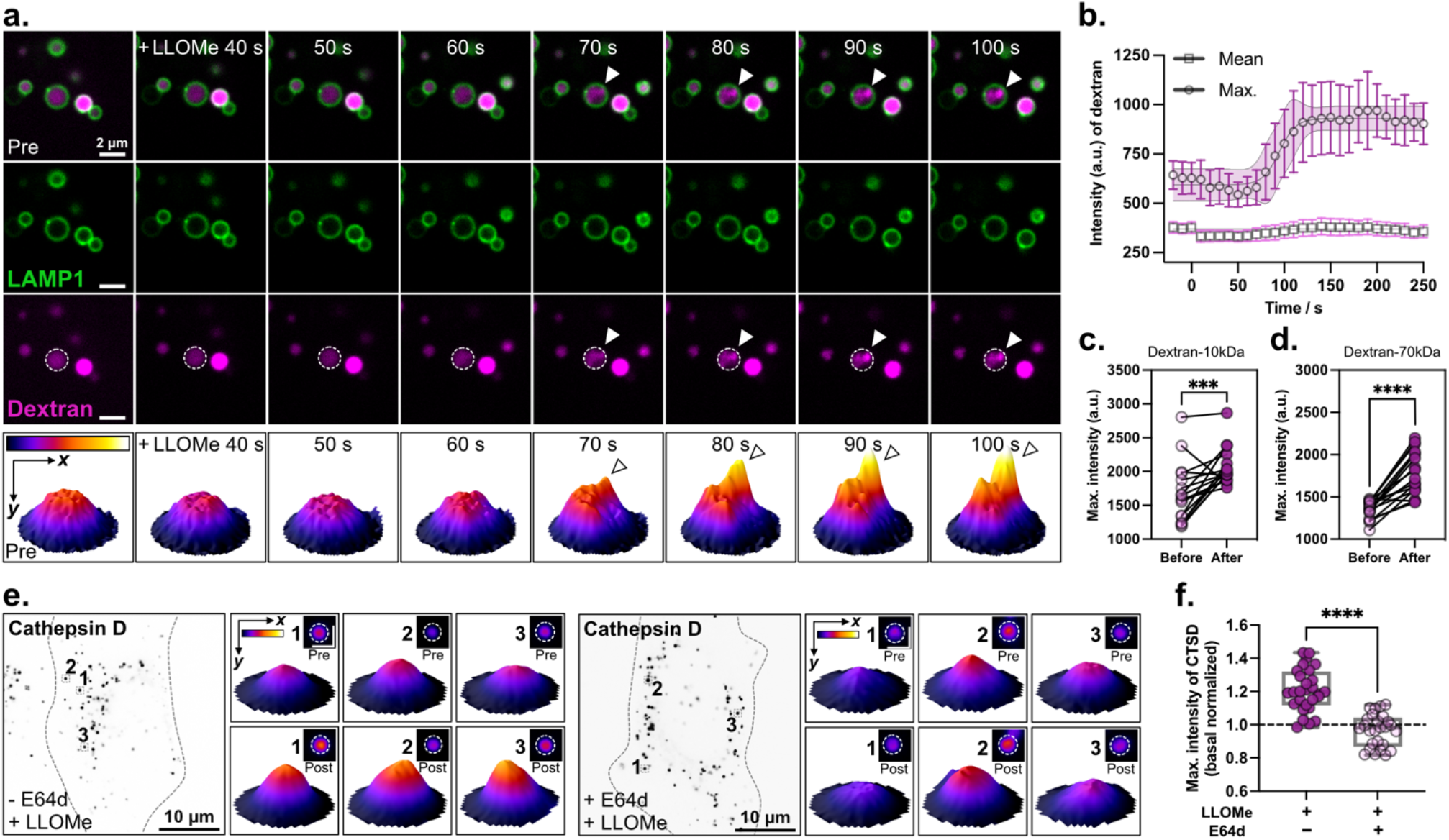
Rapid formation of intraluminal condensate upon endolysosomal damage. **a**, Representative live-cell imaging sequence of endolysosomes in U2OS cells expressing mNeonGreen-LAMP1 and loaded with 10 kDa Alexa Fluor 647-dextran, before and after LLOMe addition. Arrowheads indicate dextran condensate within LAMP1+ compartments. Bottom: 3D surface plots of fluorescence intensity for the outlined region. **b**, Maximal and mean fluorescen2ce intensity of dextran within LAMP1-outlined regions over time (10-s intervals) for condensation events as shown in **a**. Solid line, sigmoidal fit (mean ± 95% CI), over three independent experiments. **c, d**, Maximal fluorescence intensity of individual puncta of 10 kDa Alexa Fluor 647-dextran (**c**) or 70 kDa Texas Red-dextran (**d**) before and after LLOMe. Connected dots represent longitudinal measurements from the same cell (n = 17 cells over 3 independent experiments). Two-tailed paired Student’s *t*-test. **e**. U2OS cells expressing mScarlet-I-Cathepsin D pre-treated with or without cathepsin inhibitor E64d before (top) and after (bottom) LLOMe treatment. 3D surface plots of mScarlet-I-Cathepsin D fluorescence (inserted area, 1.5 µm). **f**, Normalized maximal intensity of cathepsin D puncta in cells treated with LLOMe alone (n = 28 cells) or with E64d pre-treatment (n = 26 cells), from 3 independent experiments. Two-tailed unpaired Student’s *t*-test.

We next investigated whether resident endolysosomal proteins undergo similar condensation. Consistently, the maximal fluorescence intensity of mScarlet-I–tagged cathepsin D rapidly increased following LLOMe treatment (**Fig. 1e, f**). Pretreatment with E64d abolished cathepsin D condensation (**Fig. 1e, f**), suggesting that the membranolytic activity of LLOMe, rather than its direct chemical interaction with luminal contents, is required to trigger condensation. Together, these data indicate that endomembrane damage triggers a relocalisation of intraluminal contents into condensates, prompting us to further characterise the nature of these condensates and their relationship to endolysosomal membrane damage.

### Aggregate-like protein condensates form in damaged endolysosomes

We characterised these luminal condensates with Proteostat, a rotor-based probe that becomes highly fluorescent upon immobilization within the contexts of crowded and hydrophobic protein assemblies^37,38^. Proteostat signal increased progressively over the course of LLOMe treatment (**Fig. S1a**). This increased signal was accompanied by a striking induction of Proteostat-positive puncta within LAMP1-positive endolysosomes and a significant shift of the endolysosomal content to a Proteostat-positive state (**Fig. 2a–c**), with Proteostat colocalising with condensed luminal dextran (**Fig. S1b, c**). Immunoblotting of soluble and insoluble proteins further revealed that a subset of endolysosomal proteins (*e*.*g*., CTSB, CTSD, TPP1, GBA), but not other organelle proteins, partitioned into the insoluble fraction following LLOMe-induced damage, while prosaposin (PSAP) and the endomembrane protein (LAMP1) did not, indicating selective rather than non-specific bulk aggregation of luminal contents (**Fig. 2d, e**).

**Figure 2.**
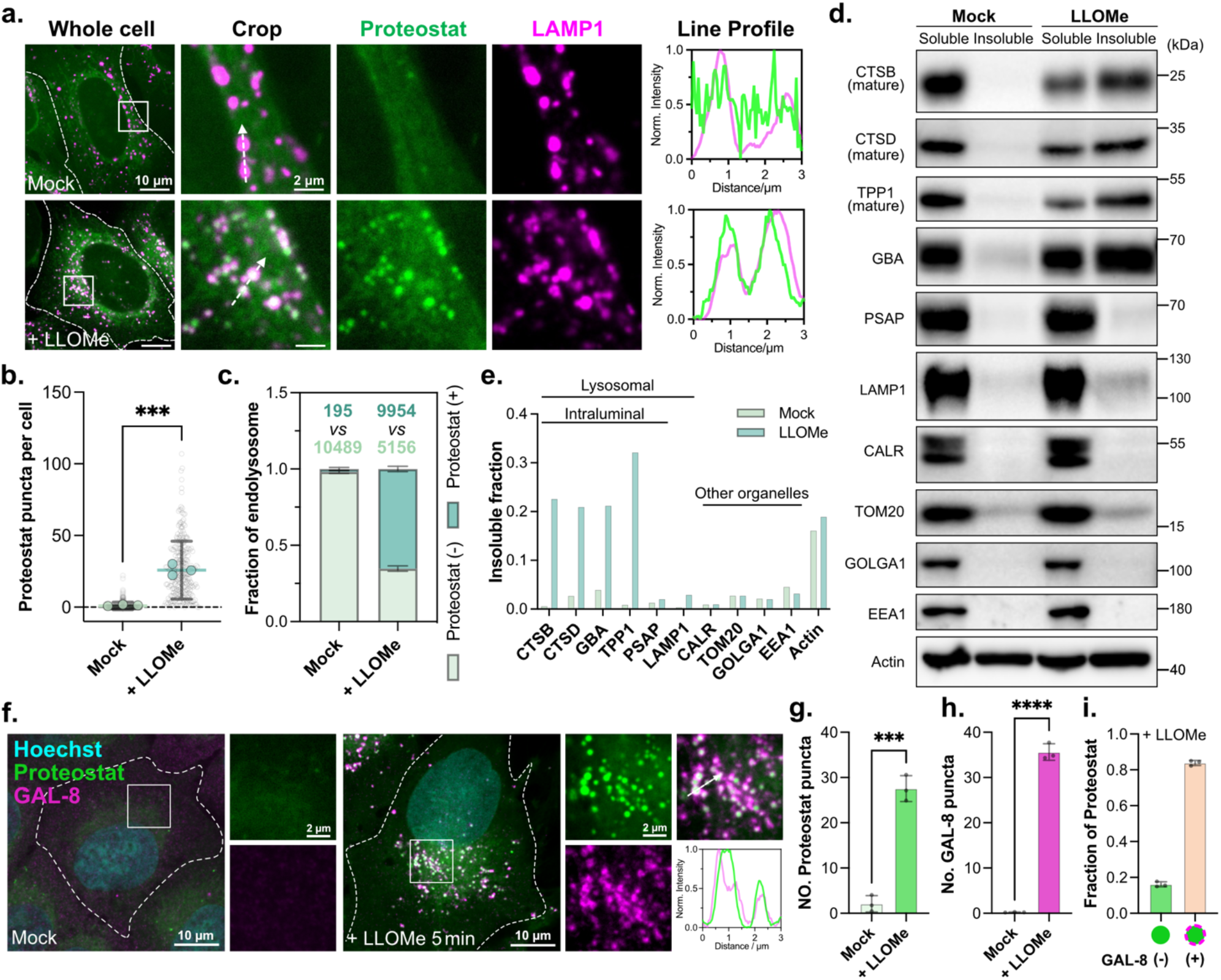
Aggregate-like protein condensates form in damaged endolysosomes. **a**, Representative immunofluorescence images of mock or LLOMe-treated U2OS cells, co-stained with Proteostat (protein aggregates) and LAMP1. Pixel intensity plot for dashed lines. **b**, Quantification of the number of Proteostat puncta per cell, for mock (n = 271 cells) and LLOMe (n = 289 cells) treated U2OS cells from 3 independent experiments, as shown in **a**. Two-tailed unpaired Student’s t-test. **c**, Quantification of the total fraction of Proteostat-positive or negative endolysosomes, for mock and LLOMe-treated U2OS cells from 3 independent experiments. **d**, Immunoblots for protein levels of soluble (RIPA supernatant) and pallet (solubilized by 4% SDS) fractions in U2OS cells (mock or LLOMe-treated for 10min). **e**, Quantification of protein insolubility in mock or LLOMe-treated cells, as immunoblotted in **d. f**, Representative immunofluorescence image of U2OS cells treated with mock (left) or LLOMe-treated for 5min (right), stained for GAL-8, Proteostat, and Hoechst. Pixel intensity plot for the dashed line. **g–i**. Graphs show number of Proteostat (**g**) and GAL-8 (**h**) puncta per cell with mock or LLOMe treatment, and the fraction of GAL-8(-) Proteostat or GAL-8(+) Proteostat per cell (**i**), after LLOMe treatment, as shown in **f**. n = 206 cells in 3 independent experiments. Two-tailed paired Student’s *t*-test.

We next co-labelled Proteostat with endomembrane damage markers to investigate whether condensate formation occurs in response to endolysosomal membrane damage. Immunostaining for endogenous Galectin-8 (GAL-8), a cytosolic lectin recruited to damaged endolysosomes after exposure of luminal glycans to the cytoplasm, revealed that over 80% of Proteostat-positive aggregates were GAL-8-positive following LLOMe treatment (**Fig. 2f–i**). This correlation extended to the ESCRT-III repair machinery: nearly 90% (a slightly higher fraction than for GAL-8) of Proteostat assemblies were positive for CHMP4B, a core ESCRT-III component potentially reflecting the reported sensitivity of the repair machinery to even minor membrane damage events (**Fig. S1d–g**).

This phenomenon was not specific for LLOMe treatment. Glycyl-L-phenylalanine 2-naphthylamide (GPN) treatment, which disrupts endolysosomal integrity by osmotic swelling^39^, also induced Proteostat-positive intraluminal assemblies colocalizing with GAL-8-marked damaged compartments in U2OS cells (**Fig. S2a, b**). Notably, E64d, which suppressed LLOMe-induced Proteostat puncta formation, failed to abolish GPN-induced condensates (**Fig. S2c–f**), indicating that cathepsin activity is dispensable under osmotic damage conditions and highlighting a distinct upstream trigger depending on the mode of membrane damage. Moreover, this effect was also observed upon silica crystal treatment, which induced Proteostat-positive condensates colocalizing with GAL-3, the other damage-sensing lectin, in THP1-derived macrophages (**Fig. S2g, h**). Altogether, these data show that intraluminal condensate formation is a general cellular response to endolysosomal injury.

### Intraluminal condensates possess a compacted ultrastructure

We then investigated the ultrastructure of these intraluminal condensates by electron microscopy. This revealed two morphologically distinct states of endolysosomal luminal contents: compartments with dispersed, granular content and compartments with electron-dense, compacted content (**Fig. 3a**). The condensed state was significantly more prevalent in LLOMe-treated cells compared to untreated controls, consistent with damage-induced condensate formation (**Fig. 3b**). Most endolysosomes having this condensed luminal morphology were tightly surrounded by endoplasmic reticulum (ER) membranes (**Fig. 3a**).

**Figure 3.**
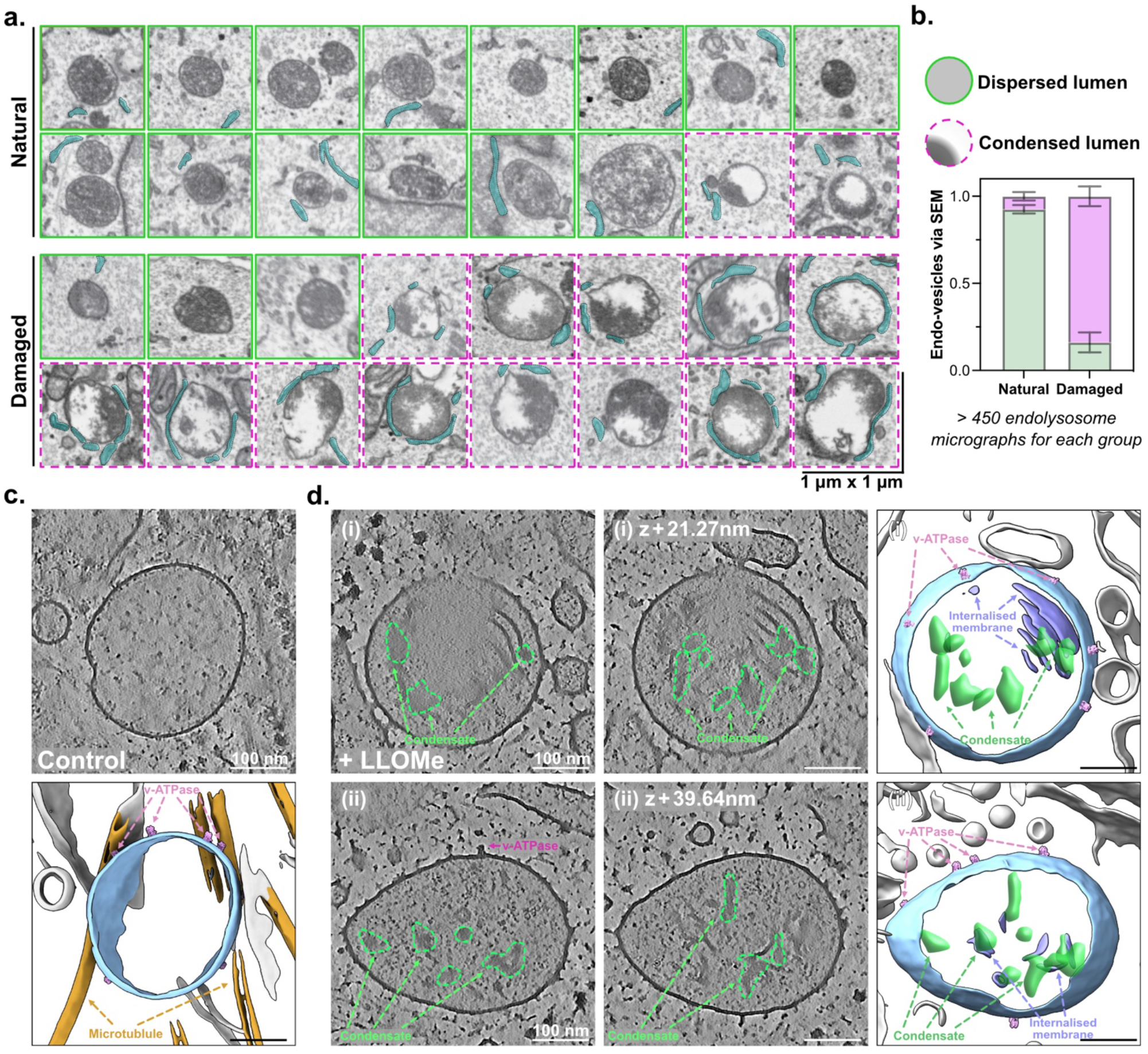
Ultrastructure of intraluminal condensates. **a**, Representative electron micrographs of endolysosomes at natural, damaged (LLOMe treatment) states. Cyan-outlined regions indicate endoplasmic reticulum surrounding endolysosomes. **b**, Schematic showing the endolysosome with dispersed or condensed luminal content (top) and fraction quantification (bottom) of each endolysosome category at natural and damaged states, as shown in **a**. n > 450 from three independent experiments. **c, d**, Representative cryo-electron tomograms of endolysosomes from untreated control (**c**) and LLOMe-treated (**d**) U2OS cells. Shown are x–y slices from 3D tomograms at different z-positions, and corresponding segmentations. The distance between two slice is (**c**) 21.27 nm and (**d**) 39.64 nm. Control endolysosomes lack discernible intraluminal condensates, whereas LLOMe-treated endolysosomes display prominent intraluminal condensates (outlined in green), clearly distinguishable from internalized membrane structures. In the segmentations, endolysosomal membranes are shown in blue, microtubules in brown, other cellular membranes in grey, V-ATPase in pink, internalized membranes in purple, and condensates in green. Tomographic slice thickness, 48.3 Å.

To characterise the architecture of intraluminal condensates in their native state, we performed cryo-electron tomography (cryo-ET), avoiding potential chemical fixation and staining artefacts occurring with conventional room-temperature electron microscopy. Endolysosomes were identified by the presence of v-ATPase-like complexes on the membrane, surrounded by the ER tubules and sheets^40^. Consistent with conventional SEM observations of electron-dense clustered structures, LLOMe-treated endolysosomes exhibited prominent intraluminal condensates, which were absent in untreated controls (**Fig. 3c, d**). Within the lumen, these condensates were largely disconnected from the membrane. While mostly lacking defined structures, they were clearly segregated from other soluble proteins in the lumen. Their morphology was distinct from internalized membrane structures, as shown by membrane segmentation, though some displayed close spatial association with the membrane (**Fig. 3c, d**). These electron microscopic characterizations provided confirmatory evidence that LLOMe treatment triggers endolysosomal intraluminal aggregation.

### Intraluminal condensate formation and dissociation are coupled to endolysosomal pH and homeostasis

Endolysosomal membrane damage was accompanied by rapid luminal pH neutralization. LLOMe treatment induced both LysoTracker Red quenching and a concurrent increase in dextran maximal fluorescence intensity, confirming that condensate formation occurs when lysosomal pH increases (**Fig. 4a–d**). To define a possible contribution of intravesicular pH disruption without (or with less) acute membrane damage, we treated cells with BafA1 or chloroquine for 24 h to neutralize the lysosomal pH. Both treatments increased Proteostat-positive puncta (**Fig. 4e–j**). However, unlike LLOMe-induced condensates, which colocalise with GAL-8 in over 80% of cases (**Fig. 2f–i**), the majority of BafA1- and chloroquine-induced Proteostat puncta showed lower GAL-8 colocalization, with most Proteostat-positive structures forming in the absence of the damage marker (**Fig. 4e–j**). These observations suggest that loss of lysosomal acidic pH alone induces intraluminal protein condensation.

**Figure 4.**
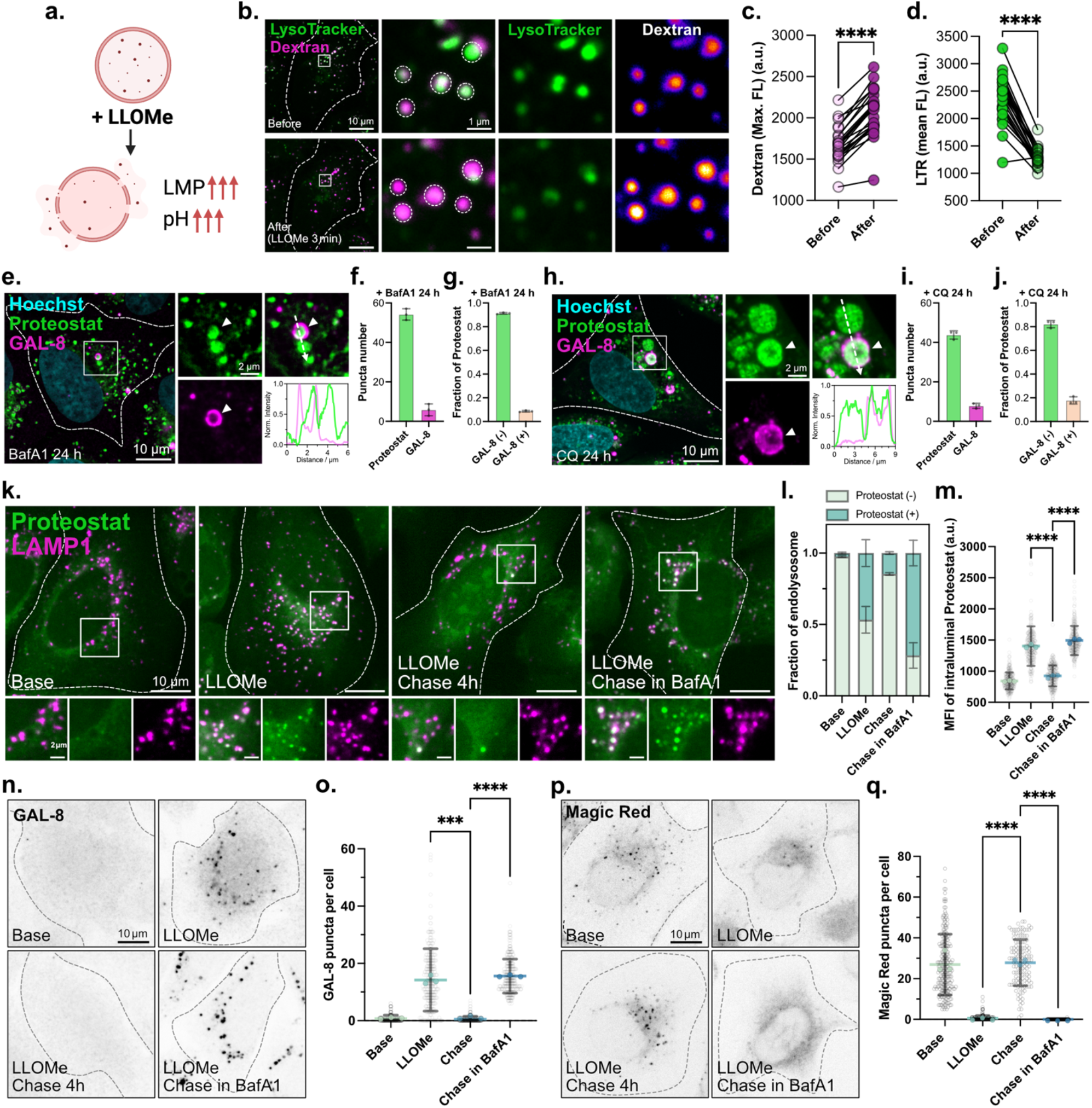
Lysosomal pH is required for the formation and dissociation of intraluminal condensates. **a**, Schematic showing LLOMe induce endolysosomal permeabilization (LMP) and pH increase. **b**, Representative images of 10-kDa Alexa Fluor 647 dextran loaded U2OS cells stained with LysoTracker Red for 30 min, before and after 3 min LLOMe treatment. **c**, Quantification of maximal fluorescence intensity of 10-kDa Alexa Fluor 647 dextran before and after 3 min of LLOMe treatment, as shown in **b**. n = 25 cells, from 3 independent experiments, Two-tailed paired Student’s t-test.. **d**, Quantification of mean fluorescence intensity of LysoTracker Red before and after 3 min of LLOMe treatment, as shown in **d**. n = 25 cells, from 3 independent experiments, Two-tailed paired Student’s *t*-test.. **e**, Representative immunofluorescence images of U2OS cells treated with BafA1 24 h, stained for GAL-8 and Proteostat. Arrowheads indicate GAL-8 positive intraluminal Proteostat puncta. Pixel intensity plot for the dashed line. **f, g**, Graphs show number of Proteostat and GAL-8 puncta per cell (**f**) and the fraction of GAL-8(-) Proteostat or GAL-8(+) Proteostat per cell (**g**), after BafA1 24 h treatment, as shown in **e**. n = 172 cells from 3 independent experiments. **h**, Representative immunofluorescence images of U2OS cells treated with chloroquine (CQ) 24h, stained for GAL-8 and Proteostat. Arrowheads indicate GAL-8 positive intraluminal Proteostat puncta. Pixel intensity plot for the dashed line. **i, j**, Graphs show number of Proteostat and GAL-8 puncta per cell (**i**) and the fraction of GAL-8^-^ Proteostat or GAL-8^+^ Proteostat per cell (**j**), after CQ 24h treatment, as shown in **h**. n = 138 cells from 3 independent experiments. **k**, Representative immunofluorescence images of U2OS cells stained for Proteostat and LAMP1. Cells were either base (untreated), treated with LLOMe 2 min, or followed by a 4-h chase in the absence or presence of 100 nM BafA1. **l**, Fraction of Proteostat(+) or (-) endolysosomes for each condition, as shown in **d**. n > 300 cells for each group, from 3 independent experiments, **m**, Quantification of mean fluorescence intensity of Proteostat in LAMP1 outlined lumen regions for each condition, n > 300 cells for each group, from 3 independent experiments, as shown in **d**. Two-tailed unpaired Student’s *t*-test. **n**, Representative immunofluorescence images of U2OS cells stained for Galectin-8. Cells were either base (untreated), treated with LLOMe for 2 min, or followed by a 4-h chase in the absence or presence of 100 nM BafA1. **o**, Quantification of GAL-8 puncta per cell for conditions as shown in **n**, n > 300 cells for each group, from 3 independent experiments. Two-tailed unpaired Student’s *t*-test. **p**, Representative images of U2OS cells incubated with Magic Red. Cells were either base (untreated), treated with LLOMe for 2 min, or followed by a 4-h chase in the absence or presence of 100 nM Bafilomycin A1. **q**, Quantification of Magic Red puncta per cell for conditions as shown in **p**, n > 130 cells for each group, from 3 independent experiments. Two-tailed unpaired Student’s *t*-test.

Having established that neutralization of lysosomal pH contributes to condensate formation, we next asked whether restoration of lysosomal homeostasis, including restoration of pH, is required for condensate dissociation. Following LLOMe washout, we monitored Proteostat, GAL-8, and the cathepsin proteolytic activity reporter Magic Red^5,22^ over time. Proteostat signal within LAMP1-positive endolysosomes increased with LLOMe treatment and dissipated during the recovery chase (**Fig. 4k–m**), coinciding with resolution of GAL-8 puncta (**Fig. 4n, o**) and re-establishment of lysosomal enzymatic activity (as judged by Magic Red signal) (**Fig. 4p, q**). When BafA1 was applied during the chase period to prevent lysosomal reacidification and function, Proteostat-positive structures persisted, coinciding with the failure to resolve GAL-8 puncta (**Fig. 4n, o**) and restore the Magic Red signal (**Fig. 4p, q**). Thus, lysosomal re-acidification is required for condensate dissociation. These findings demonstrate that lysosomal pH is critical for both intraluminal condensate formation and dissociation and that the dynamic transition of damage-induced condensates is tightly coupled to endolysosomal homeostasis.

### Oligonucleotide therapeutics are sequestered into intraluminal condensates within leaky endolysosomes

The endolysosomal lipid bilayer is perceived to be the primary intracellular barrier to endosomal escape, limiting the efficacy of oligonucleotide therapeutics^30^. Given that intraluminal condensates form rapidly upon endomembrane damage, we hypothesized that these condensates may limit escape of oligonucleotide therapeutics. LLOMe treatment enhanced ASO functional activity as measured by RT-qPCR, confirming that endomembrane damage is permissive for some degree of escape (**Fig. 5a, b**). Nevertheless, live-cell imaging revealed that despite LysoTracker Red (LTR) quenching indicative of membrane permeabilization, the mean fluorescence intensity of Cy5-ASOs in endolysosomal compartments did not significantly decrease (**Fig. 5c–e**). Consistently, the majority of ASO signal remained confined to endolysosomes even upon prolonged LLOMe treatment (**Fig. 5f**). These data reveal that breaching the barrier of endolysosomal membrane is insufficient for complete ASO release, pointing to an additional, previously unrecognised intraorganellar barrier and implicating the intraluminal condensates as a potential barrier to oligonucleotide escape.

**Figure 5.**
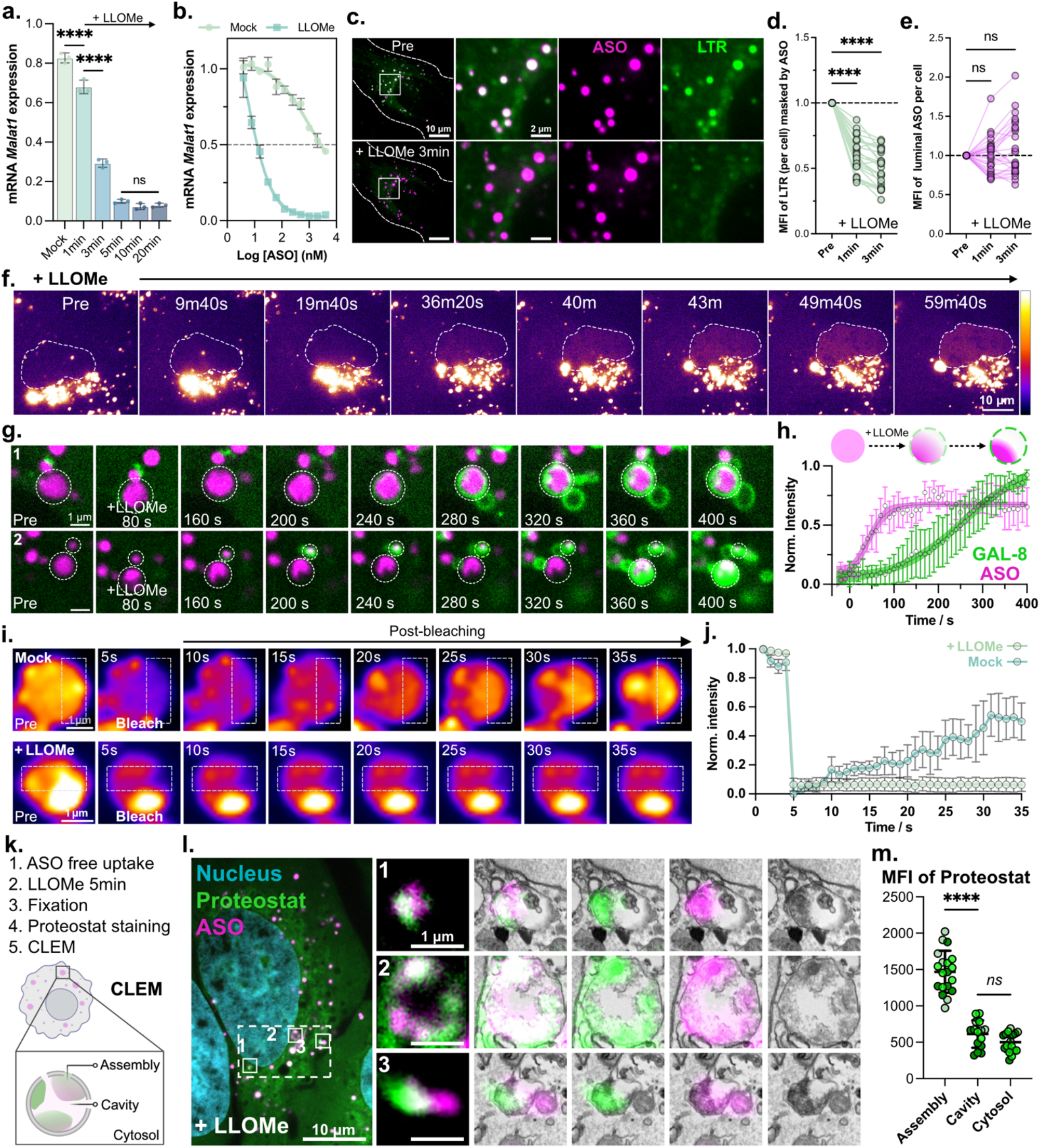
ASOs are sequestered into intraluminal condensates during endolysosomal leakage. **a**, mRNA Malat1 expression in U2OS cells treated with ASOs targeting Malat1 (100nM) for 6 h, or followed by LLOMe treatment at the indicated time, then chased up to 24 h in fresh medium, Ordinary one-way ANOVA with Tukey’s multiple comparisons test. **b**, mRNA Malat1 expression in U2OS cells treated with ASOs targeting Malat1 with various concentrations for 6 h, or followed by LLOMe treatment (3 min), then chased up to 24 h in fresh medium. **c**, Representative images of Cy5-ASO loaded U2OS cells stained with LysoTracker Red for 30 min, before and after LLOMe treatment. **d**, Quantification of mean fluorescence intensity (normalised) of LysoTracker Red before and after 1 min or 3 min of LLOMe treatment, as shown in **c**. n = 26 cells, from 3 independent experiments, Two-tailed unpaired Student’s *t*-test. **e**, Quantification of mean fluorescence intensity (normalised) of Cy5-ASO before and after 1 min or 3 min of LLOMe treatment, as shown in **c**. n = 26 cells, from 3 independent experiments, Two-tailed unpaired Student’s *t*-test. **f**, Representative live-cell imaging sequence of Cy5-ASOs endolysosomal retention during continuous of LLOMe treatment. **g**, Live-cell imaging sequence of endolysosomes in U2OS cells expressing EGFP-GAL-8 and loaded with Cy3-ASO shows ASO condensation and GAL-8 recruitment upon LLOMe treatment. **h**, Schematic and quantification of normalised ASO intensity (localised maximal) and normalised GAL-8 intensity (localised mean) over time (10-s intervals) from 13 events (3 independent experiments), as shown in **g**. Solid line, sigmoidal fit (mean ± 95% CI). **i**, Representative live-cell sequence of intraluminal ASO cargos within Apilimod-enlarged endolysosomes (top, intact mock endolysosomes; bottom, LLOMe-treated endolysosomes), as assayed by FRAP. Dashed white squares outline the bleached regions. **j**, Quantitative analysis of fluorescence intensity over time for intact mock (n = 3) and LLOMe-treated endolysosomes (n = 8), as shown in **a**, over three independent experiments. **k**, Workflow and schematic of CLEM experiments on damaged endolysosomes (treated with LLOMe 5 min) loaded with intraluminal cargo Cy5-ASOs (magenta) and stained with Proteostat (green). **l**, CLEM images show Proteostat positive and ASO positive endolysosomes (1, 2 and 3). **m**, Quantification of mean fluorescence intensity of Proteostat in regions of intraluminal assembly, intraluminal cavity and cytosol, n > 15 regions from three independent experiments, Two-tailed unpaired Student’s *t*-test.

We next investigated whether ASOs are restricted in intraluminal condensates. ASOs taken up by free uptake localise predominantly to LAMP1-marked endolysosomes (**Fig. S3a, b**). Upon LLOMe treatment, luminal ASOs relocalise from a diffuse distribution to discrete foci with increased maximal ASO intensity in LAMP1-marked compartments (**Fig. S3c, d** and **Video S5**). Consistent with our previous findings, the mean and LAMP-1 signal remained stable, mirroring the condensation behaviour of dextran and cathepsin D. A similar relocalisation was also observed upon GPN treatment (**Fig. S3e–g** and **Video S6**). ASO condensation was temporally coupled to endomembrane damage; maximal ASO fluorescence intensity increased concurrently with GAL-8 recruitment (**Fig. 5g, h** and **Video S7-9**). An equivalent temporal coupling was observed with the recruitment of the ESCRT-III component CHMP2A (**Fig. S3h, i**). To measure ASO mobility within these structures, we performed FRAP on apilimod-enlarged endolysosomes, which enabled single-endolysosome resolution while confirming that apilimod enlargement does not itself affect ASO condensation (**Fig. S3j–n** and **Video S10**). In intact endolysosomes, ASOs exhibited partial fluorescence recovery consistent with free luminal diffusion, whereas ASOs within condensate-containing damaged endolysosomes showed no fluorescence recovery (**Fig. 5i, j**), demonstrating biophysical immobilisation. The poor fluidity of condensates would prevent ASO diffusion to the cytosol even when endomembrane pores are present, providing a mechanistic basis for ASO retention within endolysosomes.

Proteostat-ASO colocalization increased significantly across distinct modes of endolysosomal damage, including LLOMe, GPN, and silica nanocrystals (compared to mock controls) (**Fig. S4a, b**). Correlative light–electron microscopy (CLEM) studies confirmed that ASOs colocalised with Proteostat-positive condensates in leaky endolysosomes, with the Proteostat signal enriched within intraluminal assemblies relative to both the luminal cavity and cytosol (**Fig. 5k–m** and **Fig. S4c**). Finally, we asked whether ASO sequestration was reversible. In apilimod-enlarged endolysosomes, ASOs transitioned from condensed foci back to a dispersed luminal distribution after recovery (**Fig. S5a**), demonstrating that sequestration is dynamically coupled to endolysosomal homeostasis. Consistently, total ASO puncta number did not change across damage and recovery conditions. However, Proteostat-ASO colocalising puncta appeared upon LLOMe treatment and dissipated upon recovery, indicating that ASOs are dynamically redistributed into and out of proteinaceous condensates without significant release from the endolysosomal compartment (**Fig. S5b–e**). These data demonstrate that intraluminal condensates represent a previously unrecognized barrier to escape of ASOs from endosomes into the cytosol, resulting in sequestration of ASOs within leaky endolysosomes.

## Discussion

Here, we describe a previously unrecognized mechanism for rapid, endolysosomal damage-induced formation of intraluminal condensates, and we show that both the assembly and dissociation of condensates are coupled to endolysosomal homeostasis. The condensate architecture revealed by cryo-ET is different from amyloid-like assemblies (**Fig. 3d**), suggesting that the endolysosomal lumen—normally a site of protein catabolism—can become a site of protein assembly under damage conditions. Whether these structures are analogous to pathological amyloids remains to be determined; however, their dissociation upon endolysosomal recovery demonstrates reversibility (**Fig. 4**), distinguishing them from irreversible aggregates. In the context of nucleic acid therapeutics, these condensates sequester luminal ASOs through biophysical immobilization (**Fig. 5i, j**), representing a newly recognized and unexpected barrier to the escape of nucleic acids from endosomes.

The molecular basis of condensate formation reflects the physicochemical sensitivity of the endolysosomal lumen, where macromolecular interactions are dependent on pH and ionic conditions. Membrane damage, whether pore-mediated (as with LLOMe) or osmotic (as with GPN), disrupts the homeostasis, for example by pH neutralization or calcium efflux, thereby altering electrostatic, hydrophobic, and hydrogen-bonding between luminal contents and thereby promoting promote condensate nucleation. Consistent with a role for pH, luminal alkalinization alone was sufficient to generate Proteostat-positive puncta, albeit with reduced colocalization with acute damage markers (compared with LLOMe-induced condensates) (**Fig. S3d**). Importantly, condensate formation appears selective for a subset of luminal hydrolases, rather than non-specific bulk aggregation of luminal contents. By sequestering soluble hydrolases within damaged compartments, intraluminal condensates may limit bystander cytotoxicity during endomembrane injury — a possibility that warrants further investigation.

The finding that intraluminal condensates sequester ASOs through biophysical immobilization has direct and clinically significant therapeutic implications. The prevailing model for endosomal escape posits that the lipid bilayers of the endomembrane system represent the primary barrier to oligonucleotide bioavailability. Yet our data reveal that even when this barrier is breached, escape remains highly restricted: condensation of luminal contents with membrane damage creates an immobilised ASO pool insulated from cytoplasmic access despite the membrane injury. The absence of FRAP recovery in condensate-associated ASOs (**Fig. 5j**) indicates effective ASO immobilization. This suggests that endosomal escape is doubly gated: first by limited leakage events from endolysosomes, and second by condensate-mediated sequestration within those compartments, a layer of restriction entirely invisible to current escape models. Notably, ASOs that reach the nucleus are sequestered within paraspeckle condensates, influencing their distribution and toxicity^24,41^, suggesting that condensate-mediated sequestration is a recurring characteristic of intracellular ASO trafficking. The reversibility of ASO sequestration (**Fig. S5**) reframes the endolysosomal compartment not as a terminal degradative sink but as a dynamic reservoir, from which cargo may be liberated by condensate dissociation. This is consistent with the prolonged pharmacodynamic activity of oligonucleotide drugs, where therapeutic effects persist for months beyond the initial phase of cellular uptake^33,42^. Targeting ASO sequestration within condensates, for example by oligonucleotide chemical modifications that reduce condensate sequestration, represents a conceptually new and unexplored strategy to improve endosomal escape and reduce the doses of oligonucleotides required for therapeutic effects.

Whether intraluminal condensates gate the cytoplasmic delivery of other nucleic acid therapeutics (*e*.*g*. mRNA) or interfere with innate immune recognition of pathogen-derived nucleic acids remains to be determined. The potential contribution of these condensates to protein aggregate nucleation in neurodegenerative disease and to lysosomal dysfunction in storage disorders remain important open questions. The identification of intraluminal condensates as a regulated, reversible organellar stress response opens a new dimension in endolysosomal biology with broad implications for cellular homeostasis and therapeutic intervention.

## Acknowledgement

This work was supported by the Hong Kong Research Grants Council General Research Fund (17102722, 17300523, 17302324) and the National Natural Science Foundation of China (32271445) to H.J. This work was supported by the Francis Crick Institute (to M.G.G.), which receives its core funding from Cancer Research UK (CC2081), the UK Medical Research Council (CC2081), the Wellcome Trust (CC2081). We thank the Imaging and Flow Cytometry Core, Center for PanorOmic Sciences, HKU, for imaging support. The work was conducted at the JC STEM Lab of Molecular Imaging, funded by The Hong Kong Jockey Club Charities Trust. For the purpose of Open Access, the author has applied a CC BY public copyright licence to any Author Accepted Manuscript version arising from this submission.

## Author contribution

H.J., M.G.G, and X.L. conceived the project. H.J. supervised the study. X.L. performed experiments and formal analysis. Y.Z. and T.N. performed cryo-electron tomography experiments and analysis. M.N., S.G.Y., C.F.B., and P.P.S. synthesized modified oligonucleotide therapeutics and designed experimental strategies for oligonucleotide studies. X.L. and H.J. wrote the original manuscript. All authors reviewed and edited the final manuscript.

## Data availability

The data underlying this article will be shared on reasonable request to the corresponding author.

## Supplementary figures

**Figure S 1.**
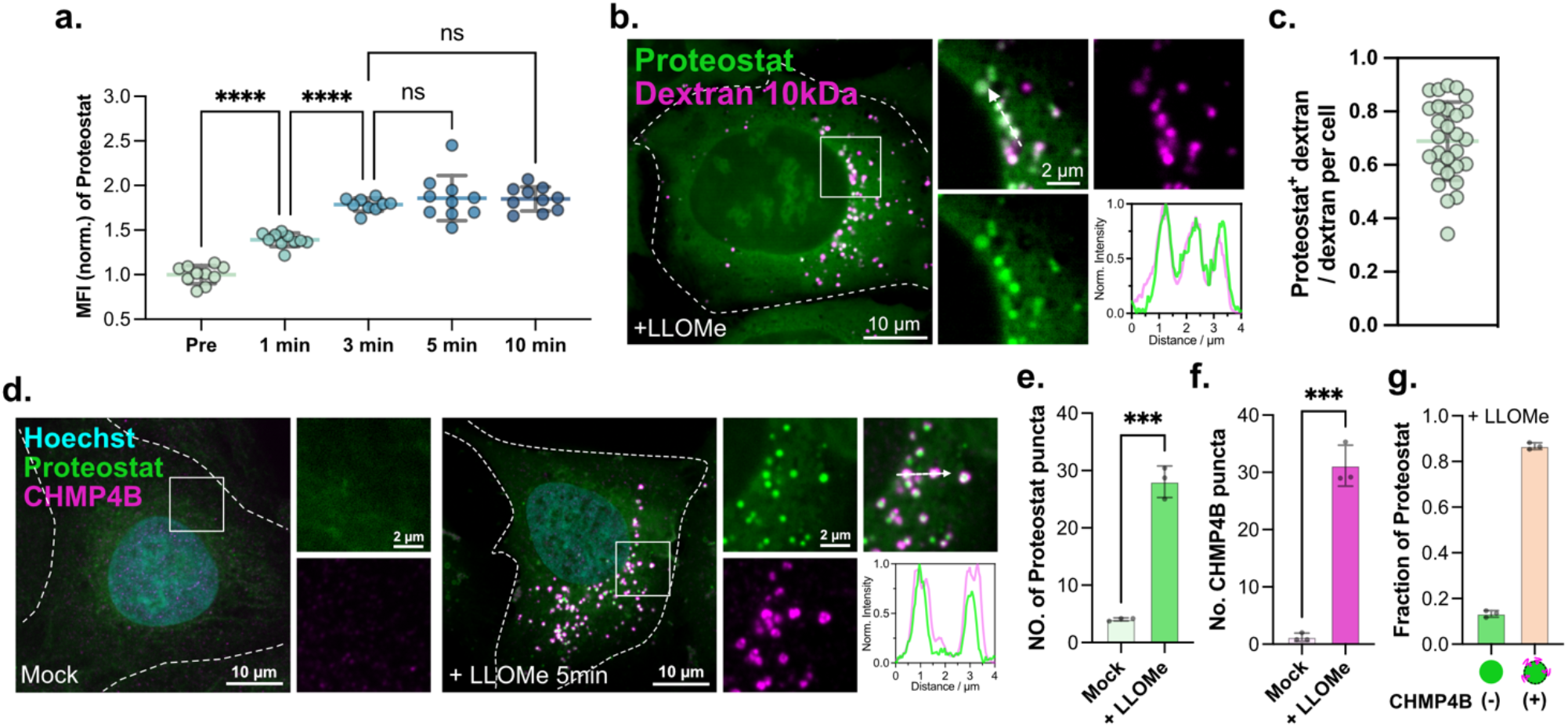
Kinetics and damage association of LLOMe-induced intraluminal condensate formation. **a**, Quantification of mean fluorescence intensity (normalised to the average value at Pre) of Proteostat upon LLOMe treatment at indicated time, Ordinary one-way ANOVA *with* Tukey’s multiple comparisons test. **b**, Representative image of U2OS cells loaded with fixable 10-kDa Alexa Fluor 647 dextran upon 3 min LLOMe treatment, stained for Proteostat. Pixel intensity plot for the dashed line. **c**, Quantification of the fraction of Proteostat^+^ dextran upon LLOMe treatment, as shown in **b**. n = 28 cells from 3 independent experiments. **d**, Representative immunofluorescence image of U2OS cells treated with mock or LLOMe 5 min, stained for CHMP4B, Proteostat, and Hoechst. Pixel intensity plot for the dashed line. **e–g**, Graphs show the number of Proteostat (**e**) and CHMP4B (**f**) puncta per cell for mock or LLOMe treatment, and the fraction of CHMP4B(+) Proteostat or CHMP4B(-) Proteostat per cell (**g**), after LLOMe treatment, as shown in **d**. n > 200 cells from 3 independent experiments. Two-tailed paired Student’s *t*-test.

**Figure S 2.**
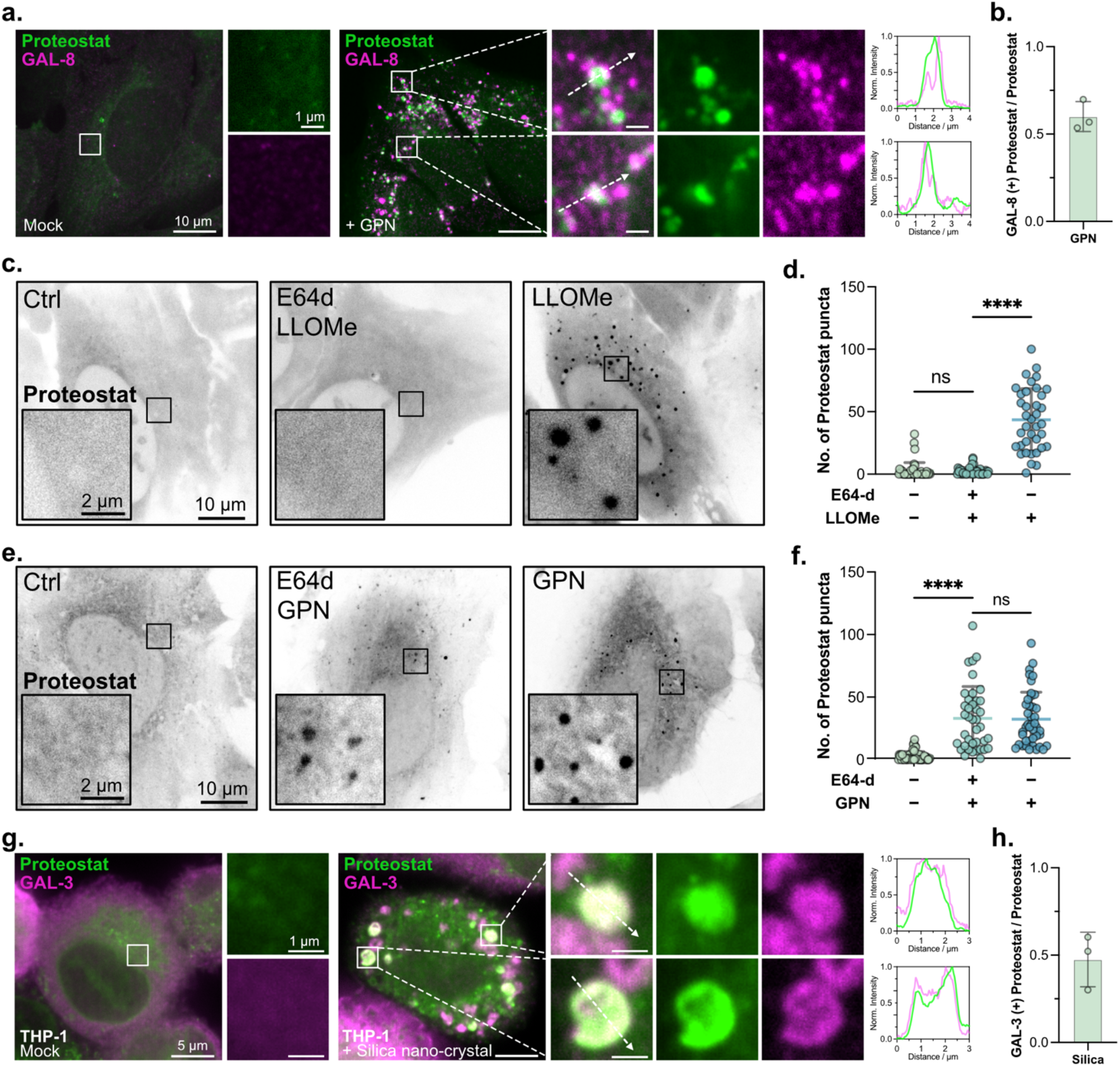
Intraluminal condensate formation is induced across different damage methods and cell types. **a**, Representative immunofluorescence image of U2OS cells treated with mock or GPN 5 min, stained for GAL-8 and Proteostat. Pixel intensity plot for the dashed line. **b**, Quantification of the fraction of GAL-8(+) Proteostat upon GPN treatment, as shown in **a**. n > 30 cells from 3 independent experiments. **c**, Representative images of U2OS cells for untreated (ctrl), E64d pre-incubation followed by treatment with LLOMe in the continued presence of E64d, and LLOMe treatment alone. **d**, Quantification of Proteostat puncta per cell for each condition, as shown in **c**. n > 30 cells for each group from 3 independent experiments, Two-tailed unpaired Student’s *t*-test. **e**, Representative images of U2OS cells for untreated (ctrl), E64d pre-incubation followed by treatment with GPN in the continued presence of E64d, and GPN treatment alone. n > 30 cells for each group from 3 independent experiments, Two-tailed unpaired Student’s *t*-test. **f**, Quantification of Proteostat puncta per cell for each condition, as shown in **e. g**, Representative immunofluorescence image of THP-1 derived macrophage cells treated with mock or silica nano-crystal for 3 h, stained for GAL-3 and Proteostat. Pixel intensity plot for the dashed line. **h**, Quantification of the fraction of GAL-3(+) Proteostat upon silica nano-crystal incubation, as shown in **g**. n > 30 cells from 3 independent experiments.

**Figure S 3.**
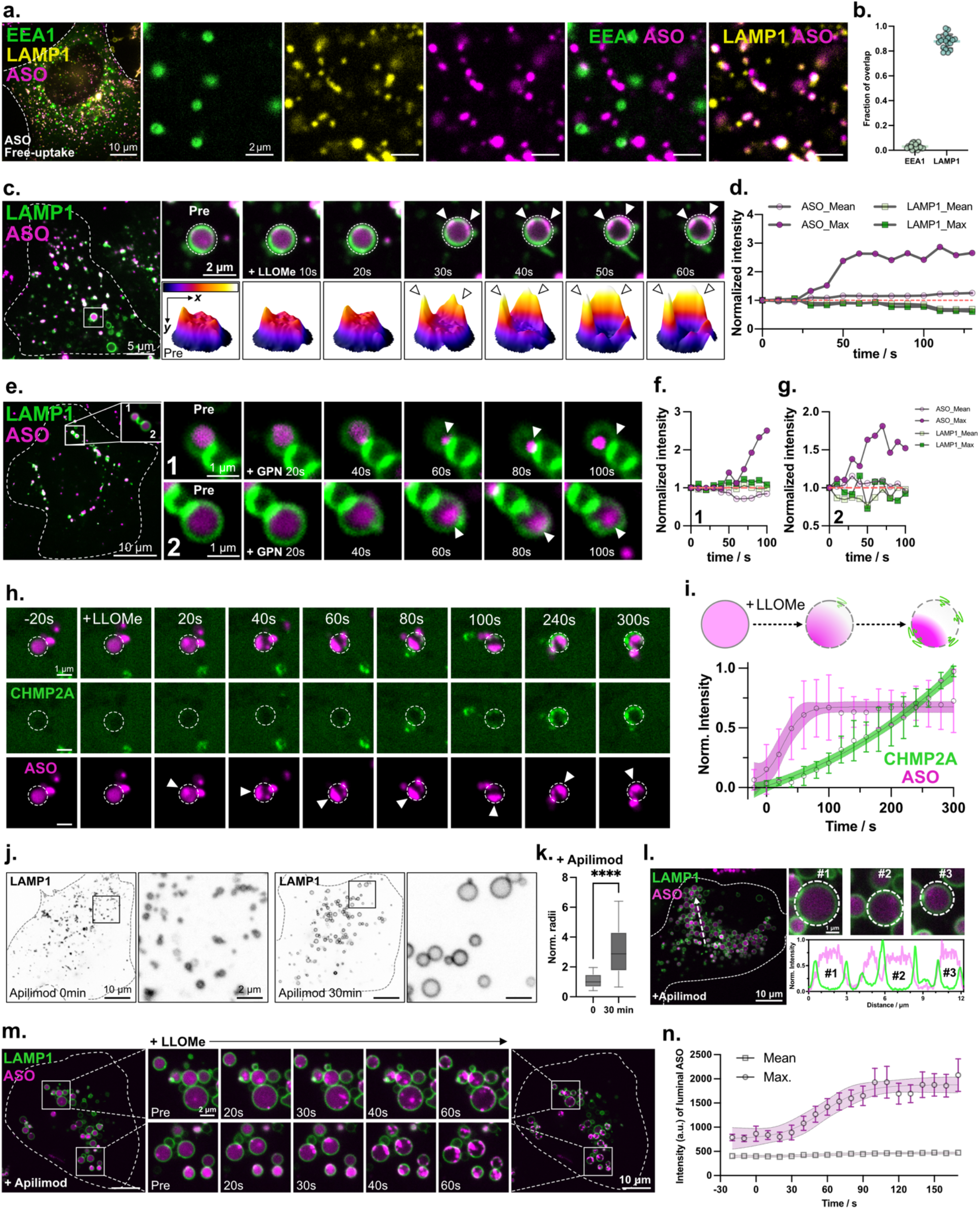
ASO condensation dynamics and characterization of ASO-containing intraluminal condensates. **a**, Representative images of U2OS cell expressing mScarlet-I-LAMP1 and EGFP-EEA1, after 6 h of Cy5-ASO free uptake. **b**, Quantification for the overlap fraction of Cy5-ASO with LAMP1 and EEA1 marked endo-vesicles, as shown in **a**. n = 19 cells from 3 independent experiments. **c**, Representative live-cell imaging sequence of endolysosomes in U2OS cell expressing mNeoGreen-LAMP1 and loaded with Cy3-ASO, treated with LLOMe. Arrowheads indicate ASO condensates within LAMP1 labelled compartment. 3D surface plots of fluorescence intensity for the outlined region. **d**, Quantification of the localised mean and maximal fluorescence of Cy3-ASO and LAMP1 within the LAMP1 outlined region over time (10-s intervals) for **a. e**, Representative live-cell imaging sequence of endolysosomes in U2OS cell expressing mNeonGreen-LAMP1 and loaded with Cy3-ASO, treated with GPN. Arrowheads indicate ASO condensates within LAMP1 labelled compartment. **f, g**, Quantification of the localised mean and maximal fluorescence of Cy3-ASO and LAMP1 within the LAMP1 outlined region over time (10-s intervals) for **e. h**, Live-cell imaging sequence of endolysosomes in U2OS cells expressing EGFP-CHMP2A and loaded with Cy3-ASO shows ASO condensation and CHMP2A recruitment upon LLOMe treatment. Arrowheads indicate ASO condensation in the dash-line outlined endolysosome. **i**, Schematic and quantification of normalised ASO intensity (localised maximal) and normalised CHMP2A intensity (localised mean) over time (10-s intervals) for 7 events from 3 independent experiments, as shown in **h**. Solid line, sigmoidal fit (mean ± 95% CI). **j**, Representative images of U2OS cells expressing mNeonGreen-LAMP1 before and after 30min of Apilimod 100nM treatment. **k**, Quantification of normalised endolysosomal radii upon Apilimod treatment, as shown in **j**. Two-tailed unpaired Student’s *t*-test. **l**, Representative image of U2OS cells expressing mNeonGreen-LAMP1 and loaded with Cy5-ASO. Pixel intensity plot for the dashed line. **m**, Live-cell imaging sequence of U2OS cells expressing mNeonGreen-LAMP1 and loaded with Cy5 conjugated ASOs, pretreated with Apilimod to enlarge endolysosomes followed by LLOMe addition. **n**, Maximal and mean fluorescence intensity of Cy5-ASOs within LAMP1-outlined regions over time (10-s intervals) to indicate the formation of intraluminal ASO condensates (n = 10 events from 3 independent experiments), as shown in **m**. Solid line, sigmoidal fit (mean ± 95% CI).

**Figure S 4.**
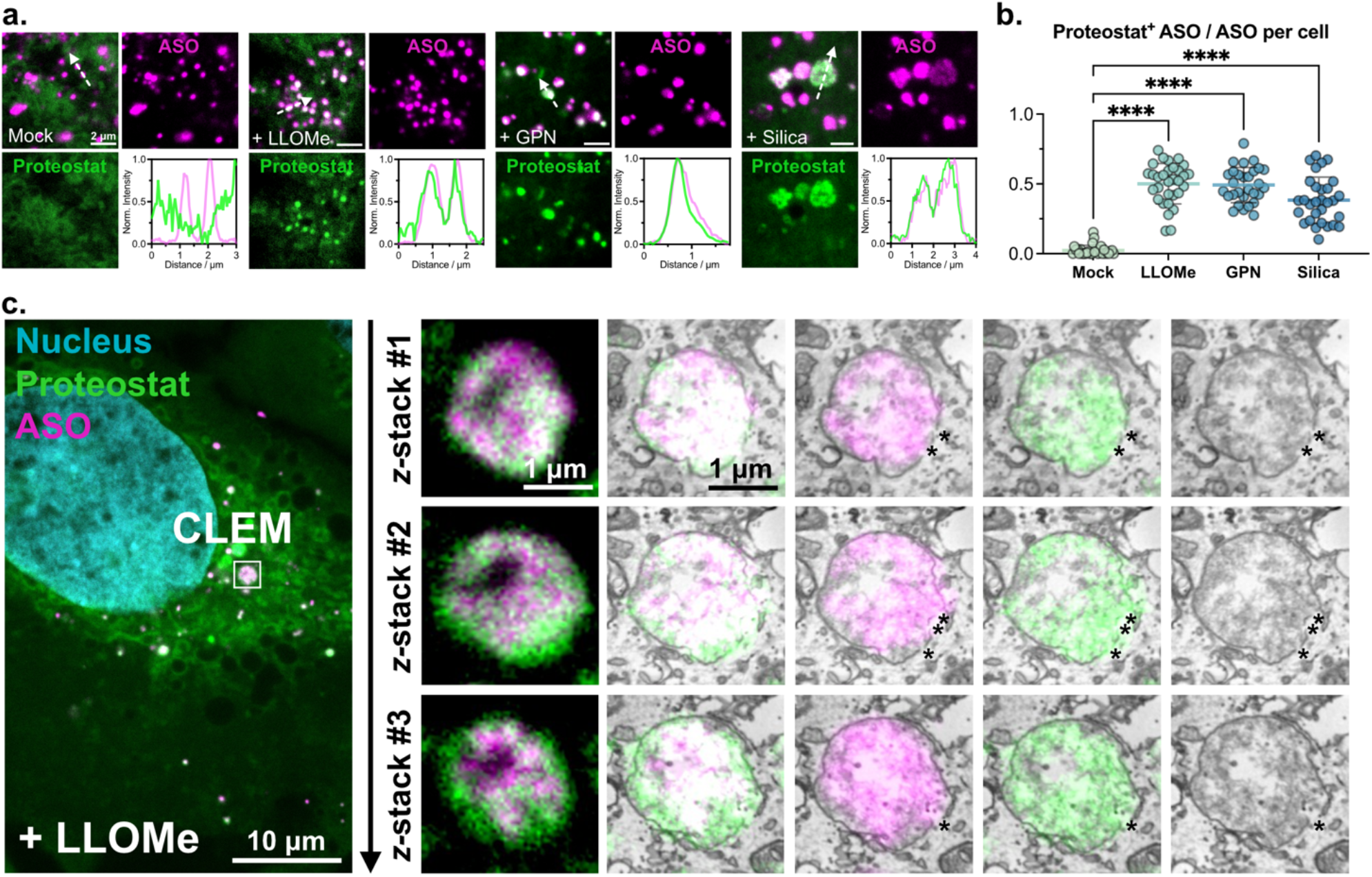
ASOs are sequestered into proteinaceous condensates in damaged endolysosomes. **a**, Representative images of U2OS cells loaded with Cy5-ASO treated with mock, LLOMe 3 min, GPN 3 min and Silica 3 h, stained with Proteostat. Pixel intensity plot for the dashed line. **b**, Quantification for the fraction of Proteostat^+^ ASO per cell upon indicated treatment, as shown in **a**. n > 28 cells for each group from 3 independent experiments. Two-tailed unpaired Student’s *t*-test. **c**, Representative CLEM image sequence shows the corresponding fluorescence–electron microscopy image overlap of damaged endolysosomes with Proteostat positive and ASO positive in z (serial stack). Asterisks indicate ASO-Proteostat condensates surrounding membrane disruption sites.

**Figure S 5.**
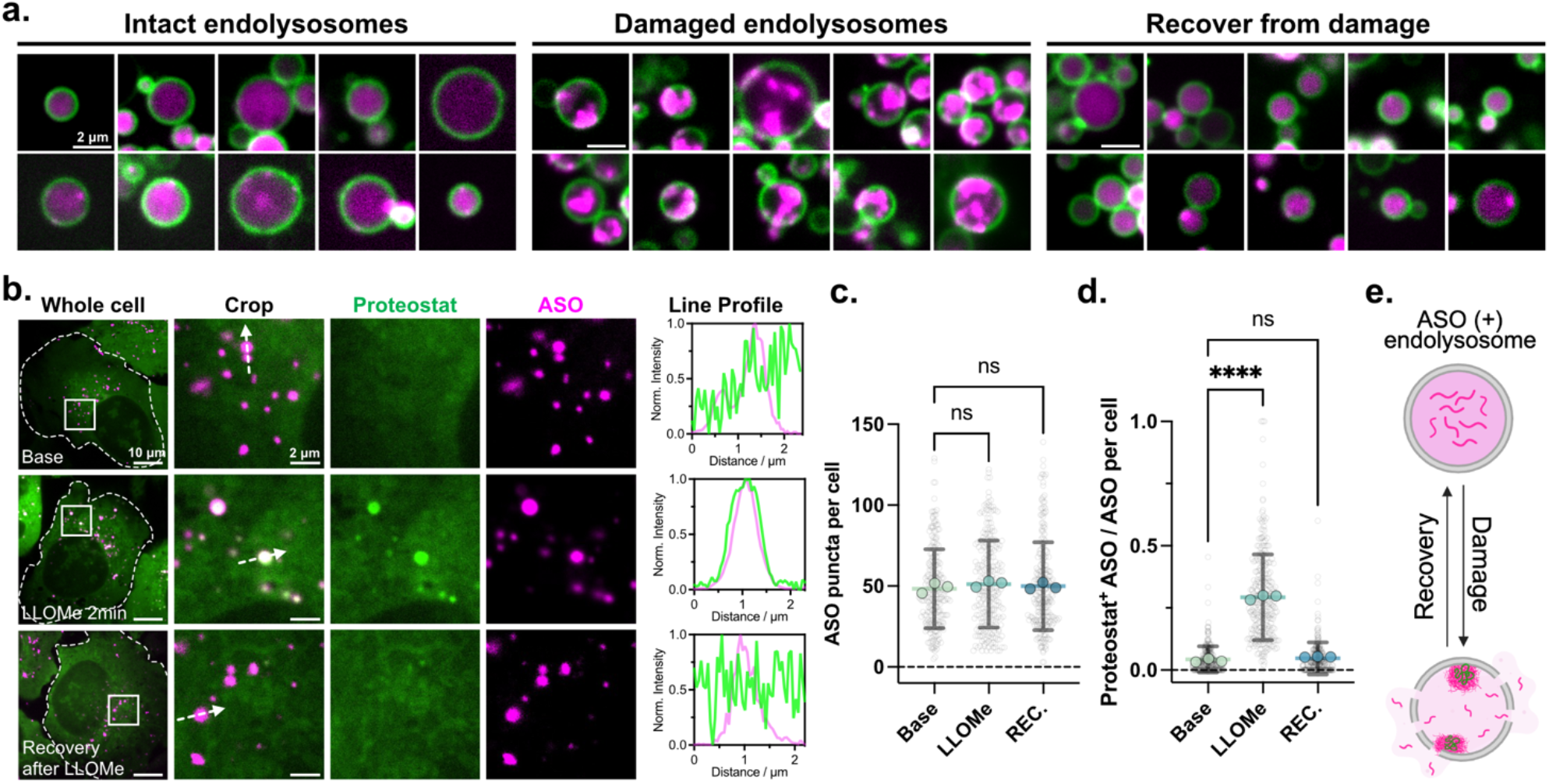
ASO sequestration is reversible upon recovery of endolysosomal homeostasis. **a**, Representative micro-fluorescence images of apilimod enlarged endolysosomal lumen loaded with Cy5-ASOs, for intact (left), damaged (middle) and recovered (right) endolysosomes, from over three independent experiments. **b**, Representative images of U2OS cells loaded with endolysosomal ASOs, stained with Proteostat at base, LLOMe 2 min and recovery after LLOMe stages. Pixel intensity plot for dashed lines. **c, d**, Quantification of ASO puncta number per cell (**c**) and percentage of Proteostat^+^ ASO per cell (**d**) at three states of endolysosomes, as shown in **b**. n > 250 cells for each state from three independent experiments. Two-tailed unpaired Student’s *t*-test. **e**, Schematic illustrating fluorescence images of reversible transition between two distinct intraluminal patterns of ASOs (magenta) and proteins (green) for intact and damaged endolysosomes, as shown in **a** and **b**.

