## Supplementary material for "Endolysosomal damage promotes intraluminal protein condensate formation that limits cytosolic leakage": Methods and Materials

### Materials and Methods

#### Cell culture and reagents

U2OS human osteosarcoma cells (American Type Culture Collection, HTB-96) were maintained at 37°C with 5% $CO_2$ in McCoy's 5A Modified Medium (Gibco) supplemented with 10% v/v fetal bovine serum (FBS) and 1% penicillin/streptomycin. Mycoplasma testing was performed monthly.

THP-1 human monocytic cell line (European Collection of Authenticated Cell Cultures, no. 88081201) were grown in suspension in RPMI 1640 medium (Gibco) supplemented with 10% v/v FBS at 37°C with 5% $CO_2$. Prior to experiments using THP-1 derived macrophages, THP-1 monocytes were seeded and cultured in µ-Slide 8-Well High glass bottom coverslips (80806, IBIDI) in RPMI with 10% FBS containing 100 nM phorbol 12-myristate 13-acetate (PMA; Sigma-Aldrich, 16561-29-8) for 24 h. After removing PMA, THP-1 macrophages were cultured in growth medium for 48 h before applied for following experiments. Mycoplasma testing was performed monthly.

Compounds were used at the following concentrations unless explicitly stated: 1 mM L-leucyl-L-leucine methyl ester (LLOMe; No. 16008, Cayman Chemical); 200 µM glycyl-L-phenylalanine 2-naphthylamide (GPN; No. 14634, Cayman Chemical); 200 µM E64d (No. 13533; Cayman Chemical); 100 nM Bafilomycin A1 (BafA1; HY-100558, MedChemExpress); 1 µM chloroquine (CQ; No. 14194, Cayman Chemical); 100 nM apilimod (HY-14644, MedChemExpress). Concentrated stock solutions of all compounds were prepared in dimethyl sulfoxide (DMSO) and stored at −80°C in single-use aliquots unless otherwise stated.

#### Antisense oligonucleotides treatment

ASOs, or Cy3 or Cy5 fluorophores conjugated ASOs as previously described[24], targeting human Malat1 were modified with a phosphorothioate (PS) backbone and 2′-constrained ethyl (cEt) wings. For imaging experiments, Cy3 and Cy5 fluorophores were conjugated to the 5′ end of the ASOs via a phosphorothioate linkage. ASOs were prepared as 1 mM stock solutions in PBS and stored at −80°C in single-use aliquots. For functional activity assays, cells at approximately 60% confluency were treated with ASOs via free uptake at the indicated concentrations and durations in complete growth medium at 37°C with 5% $CO_2$. For fluorescence imaging experiments, cells were incubated with 1 µM Cy3-ASO or Cy5-ASO in growth medium for 6 h at 37°C, followed by incubation in drug-free medium for approximately 1 hour prior to subsequent treatments to allow cellular trafficking.

#### Silica nanocrystal treatment

Silica nanocrystals (200-300 nm diameter, S131647, Aladdin) were prepared as a stock suspension in ultrapure water and then diluted in complete growth medium at 200 µg/ml, prepared immediately prior to use, and added to cells for 3 h at 37°C with 5% $CO_2$ to induce endolysosomal damage, before being fixed and prepared for immunofluorescence.

#### Drug treatment and immunofluorescence

Cells were seeded onto µ-Slide 8-Well High glass bottom coverslips (no. 80806, IBIDI) in complete growth medium under conditions appropriate for each cell line as described above. For drug treatment experiments, compounds were applied at concentrations and durations as indicated. For pulse-chase experiments, drugs were applied for the indicated duration, followed by gentle resin three times with fresh medium, and cells were then incubated in drug-free complete growth medium for the indicated chase period prior to processing.

Cells were washed three times with PBS and fixed in 4% paraformaldehyde (PFA) in PBS for 30 min at room temperature, or for 1 hour when ASO was involved. Cells were then rinsed with PBS and permeabilized in 0.2% v/v Triton X-100 in PBS for 15 min. After permeabilization, cells were blocked with 5% BSA in PBS for 30 min. Primary antibodies were diluted 1:200 in 1% BSA in PBS and incubated with cells for 1 hour at room temperature or overnight at 4°C. Cells were subsequently washed three times for 10 min each with PBS, and secondary antibodies diluted in 1% BSA in PBS were incubated for 1 hour at room temperature. Following three further 10 min PBS washes, nuclei were stained with Hoechst 33342 for 10 min and washed three times with PBS prior to imaging.

For co-immunostaining with Proteostat (ENZ-51035-KP100, Enzo Life Sciences), cells were then gently washed with PBS for 10 min after secondary antibody incubation. Then cells were stained with Proteostat diluted 1:2000 in PBS for 30 min in the dark at room temperature. Cells were washed three times with gentle rocking for 20 min, followed by Hoechst 33342 staining as described above.

The antibodies used are as follows: LAMP1 (CST, 9091), GAL3 (BioLegend, 125410), GAL8 (R&D Systems, AF1305), CHMP4B (Proteintech, 13683-1-AP).

**Plasmids**

All DNA constructs were produced using Escherichia coli DH5a (Thermo Fisher Scientific) and extracted using a plasmid midiprep kit from Qiagen. The plasmids used in this study were: mNeonGreen-LAMP1 (Addgene, 98882), mScarlet-I-LAMP1 (Addgene, 98827), EGFP-GAL-8 (Addgene, 127191), CHMP2A_GFP_N_term (Addgene, 31805), GFP-EEA1 (Addgene, 98827).

mScarlet-I-Cathepsin D was constructed by cloning the full-length coding sequence of CTSD (GenBank accession number NM_001909.5) into the LAMP1-mScarlet-I (Addgene, 98827) replacing the coding sequence of LAMP1. The full-length coding sequence of CTSD was amplified from U2OS cell cDNA library. PCR amplification was performed using PrimeSTAR GXL (Takara). The purified PCR fragment and the LAMP1-mScarlet-I vector were digested with XhoI and AgeI (NEB) at 37°C for 2 h. The digested products were ligated using DNA Ligation Kit (Takara, Cat: #6022). The primer sequences were used as follows:

CTSD Forward (XhoI): GATctcgagATGCAGCCCTCCAGCCTTC;
CTSD Reverse (AgeI): AATaccggtatGAGGCGGGCAGCCTCG.

**Cell fractionation and Western blotting**

U2OS cells with 90% confluency are cultured in 10-cm dish and replated all cells into two 10-cm dish. After 24h, U2OS cells in the 10-cm dishes were treated with LLOMe for 10 min or mock, washed three times with ice-cold PBS, and lysed in 750 µl RIPA buffer (25 mM Tris-HCl pH 7.6, 150 mM NaCl, 1% NP-40, 1% sodium deoxycholate, 0.1% SDS; no. 89900, Thermo Fisher Scientific) supplemented with Pierce Protease Inhibitor (no. A32955, Thermo Fisher Scientific). Soluble and insoluble fractions were separated by centrifugation at 18,000 × g for 15 min at 4°C. The soluble supernatant (~750 µl) was collected and combined with 250 µl 4× Laemmli SDS-PAGE sample buffer (no. 1610747, Bio-Rad) supplemented with 50 mM DTT to a final volume of 1 ml at 1× Laemmli buffer concentration. The insoluble pellet was washed three times with 500 µl RIPA buffer, with centrifugation at 18,000 × g for 15 min at 4°C between each wash, to remove residual soluble material. The washed pellet was dissolved in 300 µl PBS containing 4% SDS, 150 mM NaCl, and freshly prepared 50 mM DTT on ice, then combined with 100 µl 4× Laemmli sample buffer supplemented with 50 mM DTT to a final volume of 400 µl at 1× Laemmli buffer concentration. Both fractions were denatured by boiling at 95–100°C for 15 min and stored at −80°C until use.

To compare paired fractions from the same lysate, 5 µL of the soluble fraction and 10 µL of the insoluble fraction were loaded adjacently on the same 10% SDS-PAGE gel. The separated proteins were then transferred onto a polyvinylidene difluoride (PVDF) membrane, followed by blocking in QuickBlock Protein-Free Blocking buffer (Beyotime, P0240) for 20 min at room temperature. Membranes were then incubated with primary antibody at an appropriate dilution in primary antibody dilution solution (Beyotime, P0023A) overnight at 4ºC. After three washes in tris-buffered saline (TBS) with 0.05% Tween 20 (TBS-T) for 30 min, the membrane was incubated with species-appropriate horseradish peroxidase (HRP)–conjugated secondary antibodies for 1 hour at room temperature. Then the membrane was developed with supersensitive chemiluminescent substrate (BeyoECL Star; Beyotime, P0018AM) and imaged on a Bio-Rad ChemiDoc Imager. Antibodies used are as follows: CTSB (CST, 31718), CTSD (Proteintech, 21327-1-AP), GBA (Abcam, 309229), TPP1 (HUABIO, HA723972), PSAP (Proteintech, 10801-1-AP), LAMP1 (CST, 9091), CALR (CST, 122238), TOM20 (CST, 42406), GOLGA1 (CST, 13192), EEA1 (CST, 2411), β-actin (CST, 4967), anti-rabbit IgG-HRP (CST, 7074), anti-mouse IgG-HRP (CST, 7076).

**Super-resolution fluorescence microscopy**

Fluorescence images were acquired on a VT-iSIM super-resolution imaging system (Visitech International, UK) built around an Olympus IX83 inverted microscope (Olympus, Japan). Objectives used were a 100× oil immersion objective (NA 1.50) and a 60× silicon immersion objective (NA 1.30). Excitation was provided by a VT-LMM laser engine with 405 nm, 488 nm, 561 nm, and 640 nm laser lines. Fluorescence emission was captured simultaneously or sequentially using a VT Dual Cam image splitter and two ORCA-Quest sCMOS cameras (Hamamatsu Photonics, Japan). Z-sectioning was performed using an ASI motorized stage with piezo Z controller. Image acquisition was controlled using MicroManager (version 2.0) open-source microscopy software. For live-cell imaging, cells were maintained at 37°C with 5% $CO_2$ throughout acquisition using a stage-top incubator (Tokai Hit, Japan).

**Fluorescence recovery after photobleaching**

U2OS cells were incubated with 1µM Cy3-ASO for 6 h followed by further incubating in fresh growth medium without Cy3-ASO for 1h. After treatment of apilimod 100nM for 1 h, U2OS cells are conducted with Fluorescence recovery after photobleaching (FRAP) experiments. FRAP was performed on an imaging system using Nikon Ti2-E inverted microscope system (Nikon) equipped with Live-SR Super-resolution Unit, Yokogawa CSU-W1 Spinning Disk Unit and a bleaching module of iLAS3 Ring-FRAP/Ablation Unit. Photobleaching of Cy3-ASO was achieved using a 365-nm laser. Images were captured at 1 s intervals for at least 35 s. The photobleaching started after five time points that were used to establish the basal intensity reference.

**Antisense activities of oligonucleotides**

U2OS cells (1 × $10^5$) were seeded in 12-well plates and allowed to adhere for 16–24 h. Cells were then treated with ASOs at the indicated concentrations and durations. Following treatment, cells were washed with PBS and lysed directly in TRIzol reagent (Invitrogen, Thermo Fisher Scientific) on ice. Total RNA was extracted according to the manufacturer's instructions. Complementary DNA was synthesised using the High-Capacity cDNA Reverse Transcription Kit (Applied Biosystems, Thermo Fisher Scientific). Quantitative real-time PCR (qPCR) was performed using iTaq Universal SYBR Green Supermix (Bio-Rad Laboratories, Hercules, CA,

USA) on a Bio-Rad CFX96 Touch Real-Time PCR Detection System (Bio-Rad Laboratories). Gene expression was normalised to human Actin. The following primer sequences were used as follows:

human Malat1_forward: 5′-GAATTGCGTCATTTAAAGCCTAGTT-3′;
human Malat1_reverse: 5′-GTTTCATCCTACCACTCCCAATTAAT-3′;
human Actin_forward: 5'-ACAGAGCCTCGCCTTTGC-3';
human Actin_reverse: 5'-ATCATCCATGGTGAGCTGGC-3'.

**Electron microscopy sample preparation**

Cells with indicated treatment were processed with a protocol modified from SEM National Center for Microscopy and Imaging Research[43]. Cells were rinsed once with PBS and then once with warm 2.5% glutaraldehyde buffered in 0.1 M sodium cacodylate, then immersed in fresh fixative for 20 min at room temperature before transfer to ice for an additional hour. Buffer exchanges were performed five times (3 min per wash) using ice-cold 0.1 M sodium cacodylate, after which specimens underwent secondary fixation in 2% osmium tetroxide (0.1 M sodium cacodylate, 1 h, 4°C). Excess osmium was removed by five cold distilled water washes (5 min each), and membrane contrast was enhanced by immersion in freshly dissolved 1% thiocarbohydrazide (20 min, room temperature). Coverslips were relocated to clean wells, rinsed again five times with distilled water (5 min each), and re-exposed to 2% osmium tetroxide for 30 min at room temperature. Following five additional water washes (5 min each, cold), specimens were incubated in 2% aqueous niobium acetate at 4°C overnight. Coverslips were retrieved the next day, remove of staining by five water washes (5 min each, cold), and taken through an ascending ethanol dehydration series (30–100% in six steps, with a duplicate 100% step; 5 min each). Resin infiltration proceeded via three incubations in Embed 812 (Electron Microscopy Sciences) diluted in anhydrous acetone: 50% for 1 h, 66% overnight, and neat resin for 2 h. After blotting the reverse face to remove surplus resin, coverslips were placed cell-side down onto resin-charged BEEM capsules (Electron Microscopy Sciences) and cured at 65°C for 48 h. The blocks were cooled in liquid nitrogen (~20 seconds) and the coverslip was peeled away, leaving the embedded cells on the resin surface. The exposed cell layer was thereby embedded within the block surface. After face trimming, 100-nm sections of Ribbons were collected using a Leica UC7 ultramicrotome fitted with a Diatome diamond knife. Backscattered electron (BSE) images were acquired with a GeminiSEM 360 scanning electron microscope (ZEISS).

**Correlative light and electron microscopy (CLEM)**

U2OS cells were seeded on 35-mm gridded MatTek dishes with No. 1.5 coverslips (MatTek Corporation) and allowed to adhere for 16–24 h. Cells were then treated with 1 µM Cy5-ASO via free uptake for 6 h, followed by incubation in fresh medium for the indicated time. To induce endolysosomal membrane permeabilization, cells were treated with LLOMe for 10 min, washed twice with PBS, and fixed with 8% paraformaldehyde (PFA) in PBS for 1 hour at room temperature. After two additional PBS washes, cells were stained with Proteostat dye for 45 min at room temperature, then washed twice with PBS over 20 min. Cells were subsequently stained with Hoechst 33342 for 10 min and washed once with PBS prior to super-resolution fluorescence imaging. For correlative electron microscopy, cells were briefly rinsed with warm 2.5% glutaraldehyde buffered in 0.1 M sodium cacodylate, then immersion-fixed in the same solution overnight at 4°C before proceeding to standard electron microscopy sample preparation.

**CLEM image alignment**

Fluorescence and electron microscopy images were correlated using the BigWarp_fiji_7.0.7 plugin, which performs landmark-based deformable image registration using a thin plate spline transformation. At least 10 landmark pairs were manually placed at identifiable subcellular features visible in both modalities to constrain the transformation. The moving fluorescence image was registered to the target electron micrograph using the thin plate spline transformation. Proteostat fluorescence intensity was quantified within manually defined regions of interest corresponding to intraluminal condensate assemblies, luminal cavities, and cytosolic areas as identified in the correlated images, using Fiji. At least 15 regions from three independent experiments were analysed.

**Cryo-electron tomography (cryo-ET) and image analysis**

U2OS cells were seeded on micropatterned gold EM grids (Quantifoil R2/2, 200-mesh, Au) essentially as described previously[44]. Briefly, holey carbon–coated grids were glow-discharged for 45 s at 15 mA in a PELCO easiGlow™ system, then incubated with 1× poly-L-lysine (P8920-100ML, Sigma-Aldrich,) for 30 min. After rinsing with 0.1 M HEPES, grids were incubated for 1 h with 100 mg/ml PEG-SVA(MPEG-SVA-5000-1G, Laysan Bio), washed thoroughly with 1× PBS, and coated with PLPP gel (prepared in 70% ethanol). After drying, grids were subjected to PRIMO UV micropatterning according to the manufacturer's instructions. Patterned grids were then incubated with 20 µg/ml bovine plasma fibronectin (Sigma) in PBS for 30 min, washed three times with PBS, and exposed to UV light for 1 h. 3–4 × $10^3$ U2OS cells were seeded onto the patterned grids placed in glass-bottom confocal dishes and incubated at 37 °C with 5% $CO_2$ for 20 h until vitrification. Grids were plunge-frozen using a Leica EM GP2 (Leica Microsystems). Immediately before freezing, 1.5 µl PBS was applied to the backside of each grid, which was then back-blotted for 7 s at 20 °C and 95% relative humidity, followed by rapid plunging into liquid ethane cooled by liquid nitrogen. For LLOMe treatment, U2OS cells on grids were incubated with 1 mM LLOMe for 2 min at R.T. immediately prior to vitrification.

Clipped grids in AutoGrids (Thermo Fisher Scientific) were loaded into an Aquilos 2 cryogenic focused ion beam (cryo-FIB; Thermo Fisher Scientific) for lamella preparation. After sputter coating for 15 s and gas injection system (GIS) organometallic platinum deposition for 30 s, grids were tilted to a milling angle of 12°. Lamellae were then thinned and polished using the AutoTEM workflow, with a stepwise decrease of the gallium ion beam current from 1 nA to 30 pA, to a final thickness of ~120 nm while applying 0.2° overtilt during manual polishing. 5 lamella were prepared for each group.

Tilt series were acquired on a Titan Krios G4 transmission electron microscope (Thermo Fisher Scientific) operated at 300 kV, equipped with a Falcon 4i direct electron detector and a Selectris X energy filter (20 eV slit). Tilt series were acquired in SerialEM[45] using PACEtomo[46] at a nominal magnification of 53,000× (pixel size 2.417 Å), spanning −64° to +44° in 2° increments, with a dose of 2 $e^-/Å^2$ per tilt and a defocus range of −3 to −5 µm. The pretilt angle (-12°) was determined from defocus measurement by beam tilt[46]. Motion correction, tilt-series alignment and tomographic reconstruction were performed in AreTomo3[47] using local patch alignment. In total, 35 tomograms were reconstructed for untreated control cells and 68 tomograms for LLOMe-treated cells. All 4× binned tomograms were denoised with CryoCARE[48]. Lysosomes were identified by their characteristic ultrastructure in denoised tomograms, with this assignment supported by v-ATPase localization. Representative tomograms were further denoised and subjected to missing-wedge compensation using IsoNet2[49], followed by membrane segmentation with MemBrain-Seg[50] using default parameters. 3D visualizations, including the segmented structures and fitted V-ATPase model (EMD-44855)[40], were generated in ChimeraX (v1.10.1)[51] using the

ArtiaX plugin[52]. The V-ATPase model was placed into the potential V-ATPase density on the lysosome membrane from the denoised tomogram with an arbitrary orientation.

**Image analysis and quantification**

All fluorescence images were processed using Fiji (ImageJ version 2.14.0/1.54p). For quantification of luminal fluorescence intensity in live-cell imaging sequences, endolysosomal regions of interest (LAMP1-outlined or ASOs-covered) were delineated in Fiji by drawing a circular region of interest with a diameter matched to the size of the individual endolysosome of interest. Within each delineated region, both mean and maximal fluorescence intensities were measured at each timepoint across the imaging sequence. Three-dimensional surface plots were generated using the 'Interactive 3D Surface Plot' plugin (version V2.4) in Fiji. For puncta detection, counting, intensity measurement and object-based colocalization analysis were performed using CellProfiler (version 4.2.6). The fraction of colocalising puncta was calculated as the number of puncta in the primary channel that contained a colocalising object in the secondary channel, divided by the total number of puncta in the primary channel, and expressed as a percentage per cell.

**Statistical analysis**

Graph plots and statistical analyses were performed using GraphPad Prism 10 software (GraphPad Software, San Diego, CA, USA). Data are presented as mean ± standard deviation (SD) or mean ± 95% confidence interval (CI) as indicated in the figure legends. For comparisons between two groups, two-tailed unpaired Student's *t*-tests were used for independent samples, and Two-tailed paired Student's *t*-tests were used for longitudinal measurements from the same cell across conditions, as specified in the figure. Sample sizes (n) refer to the number of cells unless otherwise stated, and are reported in the figure legends. For sigmoidal fitting of fluorescence intensity kinetics over time, data were fit to a four-parameter logistic function and are displayed as solid lines with mean ± 95% CI. Statistical significance is indicated as follows: ns, not significant; $*p < 0.05$; $**p < 0.01$; $***p < 0.001$; $****p < 0.0001$.
